# Single-cell Transcriptomic Analysis Reveals the Cellular Heterogeneity of Mesenchymal Stem Cells

**DOI:** 10.1101/2021.11.24.469676

**Authors:** Chen Zhang, Xueshuai Han, Jingkun Liu, Lei Chen, Ying Lei, Kunying Chen, Jia Si, Tian-yi Wang, Hui Zhou, Xiaoyun Zhao, Xiaohui Zhang, Yihua An, Yueying Li, Qian-fei Wang

**Affiliations:** CAS Key Laboratory of Genomic and Precision Medicine, Collaborative Innovation Center of Genetics and Development, Beijing Institute of Genomics, Chinese Academy of Sciences, Beijing 100101, China; China National Center for Bioinformation, Beijing 100101, China; University of Chinese Academy of Sciences, Beijing 100049, China; Department of Medical Experimental Center, Qilu Hospital (Qingdao), Cheeloo College of Medicine, Shandong University, Qingdao 266035, China; International Department, Liangxiang Campus, Beijing University of Chinese Medicine, Beijing 102401, China; Yihua Biotechnology Co. Ltd, Beijing 100000, China; Peking University People’s Hospital, Peking University Institute of Hematology, Beijing 100044, China; Department of Neurosurgery, First Medical Center, General Hospital of Chinese PLA, Beijing 100853, China; Mitochondrial Medicine Laboratory, Qilu Hospital (Qingdao), Qingdao 266035, China

**Keywords:** Mesenchymal stem cell, Single-cell RNA sequencing, Heterogeneity, Lineage trajectory, Immune regulation

## Abstract

*Ex vivo*-expanded mesenchymal stem cells (MSCs) have been demonstrated to be a heterogeneous mixture of cells exhibiting varying proliferative, multipotential, and immunomodulatory capacities. However, the exact characteristics of MSCs remain largely unknown. By single-cell RNA sequencing of 61,296 MSCs derived from bone marrow and Wharton’s jelly, we revealed five distinct subpopulations. The developmental trajectory of these five MSC subpopulations were mapped, revealing a differentiation path from stem-like active proliferative cells (APCs) to multipotent progenitor cells, followed by the branching into two paths – adipogenesis or osteochondrogenesis – and subsequent differentiation into unipotent prechondrocytes. The stem-like APCs, expressing the perivascular mesodermal progenitor markers *CSPG4/MCAM/NES*, uniquely exhibited strong proliferation and stemness signatures. Remarkably, the prechondrocyte subpopulation specifically expressed immunomodulatory genes and was able to suppress activated CD3^+^ T cell proliferation *in vitro*, supporting the role of this population in immunoregulation. In summary, our analysis mapped the heterogeneous subpopulations of MSCs and identified two subpopulations with potential functions in self-renewal and immunoregulation. Our findings advance the definition of MSCs by identifying the specific functions of its heterogeneous cellular composition, allowing for more specific and effective MSC application through the purification of its functional subpopulations.

## Introduction

Mesenchymal stem cells (MSCs) are multipotent cells that can be derived from various tissues, such as adult (adipose tissue, peripheral blood, and bone marrow) and neonatal tissues (particular parts of the placenta, umbilical cord, and Wharton’s jelly) [1]. They possess self-renewal and multilineage differentiation capacities [2, 3], such as osteocytic, adipocytic, and chondrocytic differentiation. Furthermore, MSCs can secrete factors to regulate the inflammatory environment, support the development and maintenance of neurons, and promote angiogenesis and wound healing [4–6]. Due to these properties, *ex vivo*-expanded MSCs have shown promise in cellular therapy and regenerative medicine applications in recent years.

MSCs exhibit two important cellular characteristics among their properties: a high proliferation ability with differentiation potential and an immunomodulatory capability. In culture, MSCs can be expanded to produce over 10^10^ cells from an initial population of 2–5 × 10^6^ cells over 30 days of culture [7]. Even after passaging up to 10 times, MSCs are still able to maintain their proliferative and multilineage differentiation capacities, two main characteristics that define the self-renewal ability of stem cells. However, the stem cell subsets responsible for these functions have yet to be identified [8]. Another important aspect of MSCs is their immunomodulatory plasticity *via* the release of soluble factors. In particular, their therapeutic immunosuppressive capacity is mainly achieved through the production of anti-inflammatory molecules, such as prostaglandin E2 (PGE2) and TNF alpha induced protein 6 (TSG6), to inhibit the function of natural killer (NK) cells and effector T cells [4–6]. These findings suggest that MSCs may consist of a heterogeneous mixture of cells with diverse functions and multipotentiality. However, the potential cellular heterogeneity of MSCs still warrant further characterization.

*In vitro* high-capacity assays have detected tripotent, bipotent, and unipotent clones [2, 3] derived from MSCs, indicating the significant heterogeneity of MSCs in clonogenicity and multilineage differentiation. Previous studies have applied single cell RNA-sequencing (scRNA-seq) to investigate the heterogeneity of *ex vivo* cultured human MSCs. However, limited cell subpopulations were identified. Huang et al and Sun et al have highlighted that one subpopulation had strong expression of genes involved in cell cycle progression, which prevented the inference of their potential cellular functions [9, 10]. And CD142^+^ WJMSCs have been identified with wound healing potential using bioinformatic analysis [10]. In addition, many researchers only exhibited differential expressed genes and pathways after cell clustering and lack the functional assignment of the subpopulations [11, 12]. Other studies simply performed gene expression comparisons between MSCs by scRNA-seq, for example, from different sources, such as comparisons of WJ and BM, umbilical cord and synovial fluid, adipose and BM, and old and young BM [13], as well as from different stimulations, like interferon (IFN)-γ and tumor necrosis factors (TNF)-α [14]. Thus, the cellular heterogeneity associated with the proliferation, multipotency, and immunomodulatory capabilities as well as the differentiation trajectories of MSCs remains largely unclear. Biomarkers related to the enrichment of specific cells within the MSC population are also scarce.

To comprehensively investigate the cellular heterogeneity of MSCs, we profiled the single-cell transcriptome of bone marrow-derived MSCs (BMMSCs) and Wharton’s jelly-derived MSCs (WJMSCs), two essential populations of MSCs from adult and neonatal tissues, respectively. Our data revealed that five MSC subpopulations with continuous developmental hierarchies existed among MSCs. We identified a stem-like active proliferative cell (APC) subpopulation, which exhibited a strong proliferation signature and high expression levels of the perivascular progenitor markers *CSPG4/MCAM/NES* as well as stemness signatures. The APC subpopulation was located at the apex of the differentiation trajectory. Following APCs on the trajectory was the lineage-primed multipotent mesenchymal progenitor cell (MPC) subpopulation, which simultaneously expressed markers and was enriched in pathways related to osteogenic, adipogenic, and chondrogenic lineages. Interestingly, a distinct prechondrocyte subpopulation expressed higher levels of genes encoding secreted immunomodulators and possessed greater potential to suppress activated CD3^+^ T cell proliferation, supporting the role of this subpopulation in immunoregulation. Overall, our study provides a single-cell transcriptomic blueprint of MSCs and uncovers the characteristics of stem-like, highly proliferative, multipotent, and immunoregulatory subpopulations among MSCs. These findings are helpful for advancing the definition of MSCs by identifying specific subpopulations, thereby enhancing their therapeutical potential by increasing specificity.

## Results

### Characteristics and single-cell transcriptome of BMMSCs and WJMSCs

BMMSCs and WJMSCs expanded *in vitro* at passages 6–7 were applied in our study. These cells were maintained in a stable state and were used for clinical application [15–17]. First, assays to identify and characterize MSCs were performed based on the criteria published by the Mesenchymal and Tissue Stem Cell Committee of the International Society for Cell & Gene Therapy (ISCT) [18]. The MSCs maintained their adherence to plastic when cultured under standard conditions and showed the common spindle-shaped, fibroblast-like morphology (**Figure 1,** A). In an *in vitro* differentiation system, both BMMSCs and WJMSCs could differentiate into adipocytes (Figure 1B), osteoblasts (Figure 1C) and chondrocytes (Figure 1D). In addition, the panels of positive MSC markers (CD73, CD90, CD105) and negative MSC markers (CD45, CD34, CD11b, CD19, and HLA-DR) were expressed on more than 98% and less than 1%, respectively, of the MSCs (Figure S1, A and B). However, in the same culture medium, WJMSCs had higher rates of proliferation (Figure S1C) and smaller average diameters than BMMSCs (Figure S1D), which is also in line with the results of previous studies [1, 19, 20].

**Figure 1.**
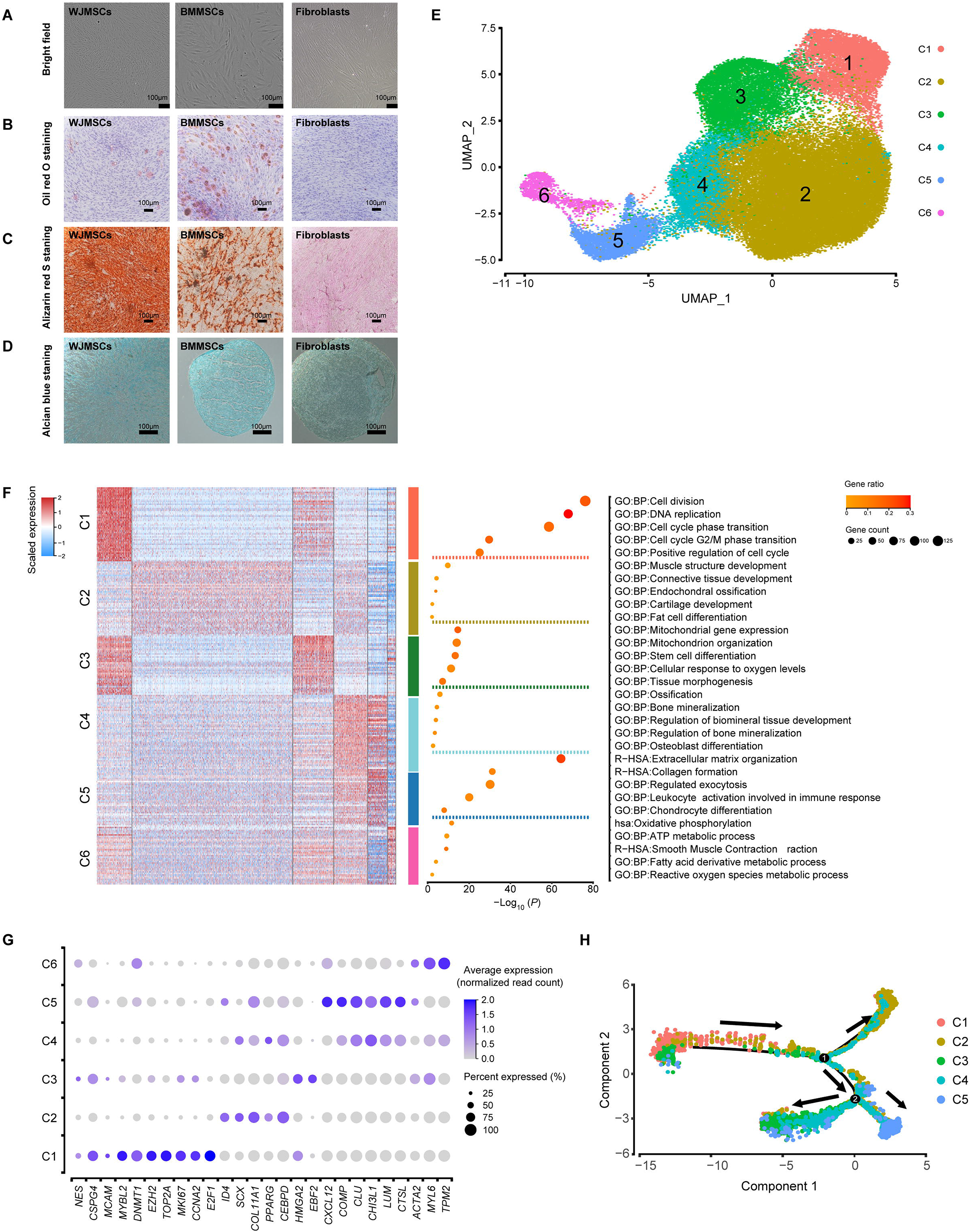
Characteristics and single-cell transcriptome profiling of BMMSCs and WJMSCs. **A.** Representative bright-field images of the WJMSCs, BMMSCs, and fibroblasts (as a control). **B.** Representative plots of WJMSCs/BMMSCs/fibroblasts stained with Oil Red O after adipogenic induction for 28 days. **C.** Representative plots of WJMSCs/BMMSCs/fibroblasts stained with Alizarin Red after osteogenic induction for 28 days. **D.** Representative plots of WJMSCs/BMMSCs/fibroblasts stained with Alcian Blue after chondrogenic induction for 28 days. **E.** Cell type identification on the UMAP plot of 33,594 cells from 3 WJMSC samples and 27,702 cells from 3 BMMSC samples. **F**. DEGs and corresponding representative GO terms. Left: Heatmap showing the significantly DEGs (expression percentage > 0.25, log2FC > 0.4) in each cluster. The color indicates the scaled expression level. Right: The enriched functional pathways in each cluster are listed. The dot size indicates the number of genes, and its color indicates the gene ratio in each cluster (gene number *versus* total gene number in the term). **G.** Dot plot showing the relative expression levels of the classical DEGs in each cluster. The dot size indicates the percentage of cells in the cluster expressing a gene; the shading indicates the relative level of expression (low to high, shown as light to dark). **H.** Pseudotime map of each subpopulation generated by Monocle. MSC, mesenchymal stem cell; WJMSC, Wharton’s jelly-derived MSC; BMMSC, bone marrow-derived MSC. C1: Cluster 1. C2: Cluster 2. C3: Cluster 3. C4: Cluster 4. C5: Cluster 5. C6: Cluster 6; UMAP, uniform manifold approximation and projection; DEG, differentially expressed genes; GO, gene ontology.

To uncover the cellular composition and diversity of MSCs, we performed scRNA-seq on 3 WJMSC and 3 BMMSC samples from different donors using the high-throughput 10x Genomics platform (Table S1). After stringent cell filtration, high-quality single-cell transcriptomes of 61,296 MSCs (33,594 cells derived from WJ, accounting for 54.8% of the total population; 27,702 cells derived from BM, accounting for 45.2% of the total population) were obtained for downstream analysis. Compared to BMMSCs, WJMSCs had higher median numbers of expressed genes (4136 vs 3144) and higher unique molecular identifier (UMI) counts (21,730 vs 13,317) (Figure S1, E and F) but a similar median percentage of mitochondrial genes (2.79% for WJMSCs vs 2.43% for BMMSCs) (Figure S1G). The average expression levels of the well-known established MSC markers CD73*/NT5E,* CD90*/THY1*, CD105*/ENG,* and *CD44* (Figure S1H) was consistently high in both WJMSCs and BMMSCs. These results indicated that we successfully obtained single-cell transcriptomes of WJMSCs and BMMSCs via a high-throughput approach for further analysis.

### Transcriptional heterogeneity exists within five distinct MSC subpopulations with unique signature

To investigate the cellular heterogeneity of MSCs, unsupervised clustering by the uniform manifold approximation and projection (UMAP) technique was performed after cell cycle regression. In total, six clusters were identified (Figure 1E and Figure S2, A to C, Table S2). Typical MSC markers, including CD73*/NT5E,* CD90*/THY1,* CD105*/ENG,* and *CD44*, showed variable expression levels among each cluster (Figure S2D), suggesting that the traditional criteria were unable to define MSC subpopulations due to their intrinsic heterogeneity. Meanwhile, based on the UMAP pattern of individual samples from these six donors, we found that BMMSCs had amore similar pattern while WJMSCs had higher individual complexity, suggesting that WJMSCs possessed higher inter-donor variability (Figure. S2E). To determine the cellular identity of each cluster, significantly differentially expressed genes (DEGs) and corresponding enriched pathways, as well as potential key regulators, were identified. Cells in cluster 1 showed a stronger characteristic of active proliferation, as these cells had high expression levels of genes related to DNA replication and cell cycle progression, including proliferation markers (*TOP2A, MKI67,* and *E2F1*) and a cell cycle regulator (*CCNA2*) (Figure 1, F and G). Furthermore, the transcription factors (TFs) E2F1 and E2F8, known cell cycle progression regulators, were predicted by SCENIC to be the active TFs in cluster 1 (Figure S2F). Interestingly, NG2*/CSPG4,* CD146*/MCAM,* and *NES* (Figure 1G), the characteristic markers of perivascular mesodermal progenitor cells [21–23], were also highly expressed in cells in cluster 1. When these observations were combined with the findings regarding the expression levels of genes essential for maintaining pluripotency and the undifferentiated stem cell state, such as *MYBL2* [24], *DNMT1,* and *EZH2* [25] (Figure 1G), cluster 1 cells were classified as potentially stem-like APCs.

Cells in cluster 2, accounting for more than half of the total cell number (Figure S2A), exhibited an expression signature enriched for trilineage differentiation, including osteogenic differentiation (*ID4* [26]), chondrogenic differentiation (*SCX* [27] and *COL11A1* [28]), and adipogenic differentiation (*PPARG* and *CEBPD*) [29] (Figure 1, F and G). The predicted TFs in cluster 2 are known to govern diverse lineage commitment decisions, including mesoderm development (*IRX3*), osteogenesis (*JUN, ATF4*) [30], chondrogenesis (*TRPS1*) [31], and adipogenesis (*CEBPB*) [29] (Figure S2E). Considering the revealed multilineage differentiation potential, cells in cluster 2 were referred to as tripotent multipotent MPCs.

In addition to stem and progenitor cells, differentiated precursors were identified. Cells in cluster 3 were enriched with genes involved in stem cell differentiation (*PSMD2, PSMD7, PHF5A*), tissue morphogenesis (*TBX3, CFL1, TRIM28*), and mitochondrial biogenesis for adipogenesis (*NDUFA9, UQCC2, ATP5F1B, COX20*) (Figure 1F). Moreover, cluster 3 cells expressed high levels of genes related to the regulation of adipocyte differentiation, such as *EBF2* and *HMGA2* (Figure 1G). GATA2, reported to be expressed in preadipocytes and to play a central role in controlling adipogenesis [32, 33], was also predicted to be the active TF in cluster 3 (Figure S2E). Thus, we referred to cluster 3 cells as unipotent preadipocytes.

Cells in Cluster 4 expressed high levels of the cartilage-specific gene *COMP* and the extracellular matrix remodeling genes *CHI3L1, CLU, LUM,* and *CTSL* (Figure 1G). Correlation analyses of the top-gene transcriptome between our analyzed MSCs and published chondrocyte and osteoblast datasets [34] revealed that cells in cluster 4, which also showed higher expressions of osteogenesis (*OMD*, *ASPN*, *GPM6B*, *IFITM1,* and *GPNMB*) (Figure S2G and H), were closely related to chondrocytes and osteoblasts. Thus, cells in cluster 4 were referred to as bipotent prechondro-osteoblasts (pre-COs). Cluster 5, on the other hand, resembled chondrocytes and had a higher expression of genes involved in chondrogenesis (*COL6A3, COL6A1,* and *ECM1*) (Figure S2G and H). Interestingly, pathways involved in immunomodulation and secretion were also enriched in cluster 5 (Figure 1F). Thus, cluster 5 cells were annotated as immunoregulatory prechondrocytes. Cells in cluster 6, accounting for the lowest proportion (2.25%) of the cell population (Figure S2A), were enriched with genes essential for smooth muscle contraction (*ACTA2, MYL6,* and *TPM2*) (Figure 1, F and G). The predicted active regulatory TFs SOX15 and NR1D2 have been demonstrated to play a key role in determining early myogenic cell development (Figure S2E) [35, 36]. Thus, cluster 6 cells were referred to as pre-smooth muscle cells (pre-SMCs), consistent with the capability of MSCs to differentiate towards vascular lineages. Cells in cluster 6 also expressed high levels of genes participating in metabolic processes (Figure 1F), supporting the importance of metabolic reprogramming during the differentiation of MSCs into SMCs [37].

To explore the relations and the developmental hierarchies among the subpopulations, we performed pseudotime analysis with Monocle2. The stem-like APCs were positioned in the “source” cell state, followed by MPCs. Then, two branching paths were derived from MPCs: one leading to pre-COs and differentiated unipotent prechondrocytes, and the other leading to preadipocytes (Figure 1H and S2I). Moreover, the Monocle2 result was supported by the RNA velocity analysis with Velocyto [38] (Figure S2J), which enables the prediction of potential directional trajectories and cell state transitions by connecting measurements to the underlying mRNA splicing kinetics. Overall, these findings revealed that MSCs were composed of heterogeneous and continually developing cell populations that progressed from stem-like APCs to tripotent MPCs and ultimately to bipotent and unipotent precursors.

### Specialized active proliferative cluster (Cluster 1) cells possess stem-like transcriptional signatures

It is generally believed that one of the key characteristics of MSCs is their ability to undergo robust proliferation ability while maintaining their multilineage differentiation potential. We performed further analysis to explore whether cluster 1 cells possessed this stem-like characteristic.

Compared with cells in the other clusters, cluster 1 cells expressed extremely high levels of NG2*/CSPG4,* CD146*/MCAM,* and *NES* (**Figure 2,** A and B). NESTIN^bright^ NG2/CSPG4^+^ periarteriolar mesodermal progenitor cells, whose marker phenotype indicates the stemness characteristics of self-renewal and differentiation into multiple mesenchymal lineages, were reported to differentiate into MSCs and to constitute the origin of MSCs in multiple organs [21, 23]. In addition, the paraxial mesoderm can bud off into MSCs in both *in vitro* and *in vivo* experiments [39–41]. To validate the identity of cluster 1, we compared the single cell transcriptome data between cluster 1 cells and NG2^+^ periarteriolar cells, LEPR^+^ perisinusoidal cells [42], and paraxial mesoderm cells, respectively [43]. We compared the overall expression pattern by performing a Pearson correlation test with genes involved in maintaining stemness, including those related to the self-renewal (*E2F8, CTCF, PBX3,* and *MYBL2*), negative regulation of cell differentiation (*ASPM, CBFB, SUZ12, WNT5A,* and *PTHLH*), and cell cycle regulation (*CDKN2C* and *CDKN1A*), etc. Clearly, cluster 1 but not cells in other clusters were closely related to NG2^+^ periarteriolar cells (*in vivo*, Figure 2C) and paraxial mesoderm cells (*in vitro*, Figure 2D and Figure S3A). Stemness genes along with those related to the notch signaling pathways (*E2F1, EZH2,* and *TFDP1*), negative regulation of apoptosis (*HMGB2, BRCA1, PAK4,* and *MAZ*) and polycomb groups (*PCGF6* and *PHC1*), were all strongly co-expressed in cluster 1 (Figure 2, C and D). These observations indicated that cluster 1 cells resembled mesodermal progenitor cells with the ability to self-renew and differentiate into multiple mesoderm lineages. Via SCENIC analysis, CTCF, EZH2, E2F8, PBX3, MYBL2, and TFDP1 were also identified as potential activated TFs in cluster 1. These TFs are transcriptional activators targeting the genes related to self-renewal pathways (Figure 2E and Figure S3B). Together, these results suggested that cluster 1 cells possessed a high proliferative capacity combined with a stem-like transcriptional signature.

**Figure 2.**
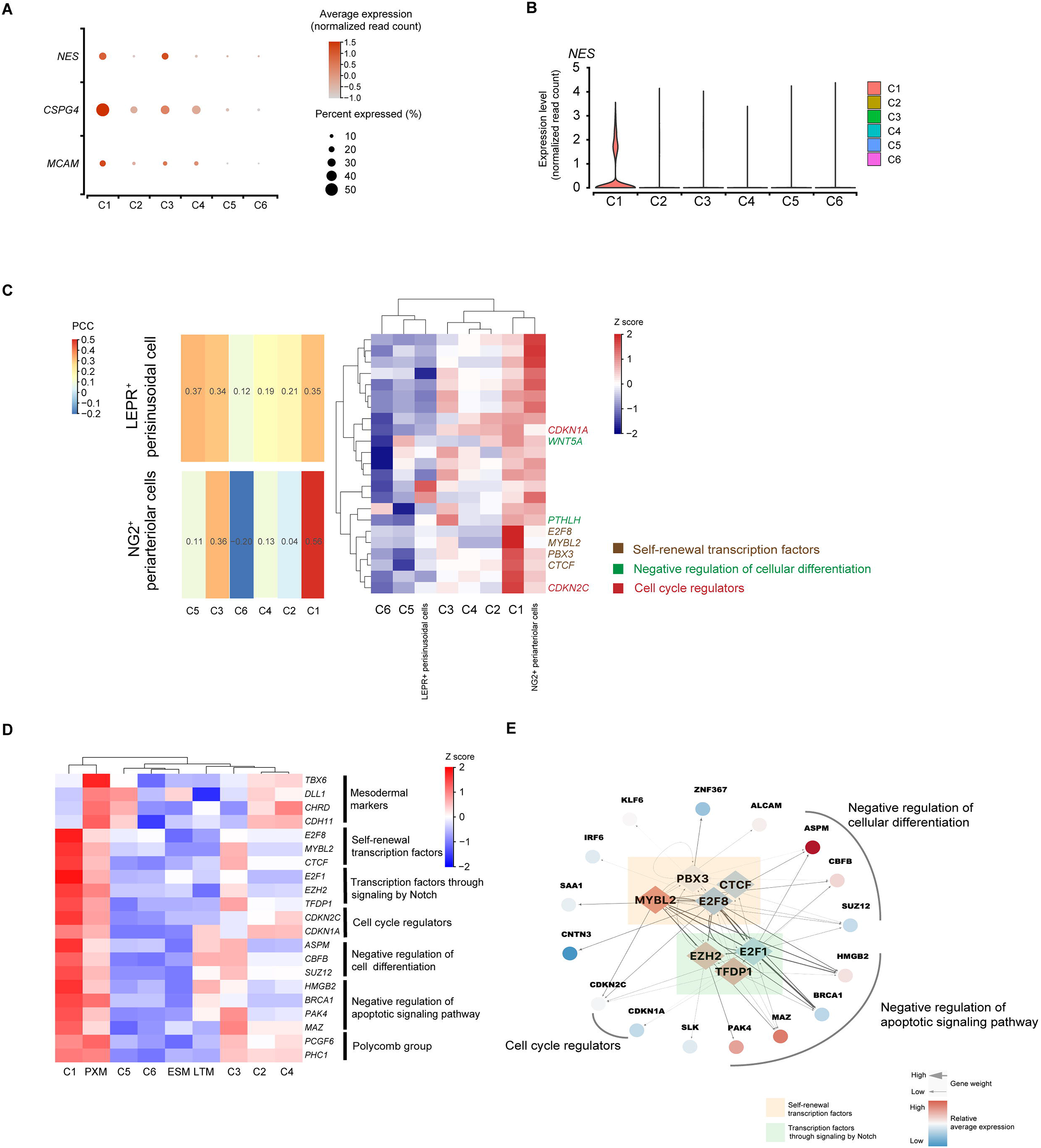
Identification of specialized stem-like cells (Cluster 1) with high proliferative capacity. **A.** Dot plot showing the relative expression levels of mesodermal progenitor cell marker genes (NG2*/CSPG4,* CD146*/MCAM,* and *NES*) in MSC subpopulations. The dot size indicates the percentage of cells in the cluster expressing a gene; the shading indicates the relative level of expression (low to high, shown as light to dark). **B.** Violin plot showing relative expression levels of mesodermal progenitor cell marker genes (NG2*/CSPG4* and *NES*) in MSC subpopulations. **C.** Data comparison between transcriptomes of MSC subpopulations and cells from published datasets. Left: Heatmap displaying the correlation matrix analyzed by PCC. Right: The expression patterns of selected genes involved in the indicated biological processes between the transcriptomes of MSC subpopulations and cells from published datasets [42], including NG2^+^ periarteriolar cells and LEPR^+^ perisinusoidal cells. **D.** Heatmap displaying the expression patterns of selected genes involved in the indicated biological processes between each subpopulation and cells from published datasets [43], including DLL1^+^ PXM, LTM and ESM. **E.** The potential activated TFs for cluster 1 predicted by SCENIC are shown as diamonds, and their potential downstream targets are shown as circles. PXM, paraxial mesoderm; LTM, lateral mesoderm; ESM, early somites; PCC, pearson correlation coefficient.

### MPCs (Cluster 2) are subgrouped into trilineage orientations

Cells in cluster 2, annotated as multipotent MPCs, exhibited expression of MSC stemness-associated markers (*CD9, CD44, ITGB1, SDC4,* and *ITGAV*) (Figure S3C). Moreover, the dot plot (Figure 1F) and pseudotime model of gene expression dynamics (Figure 3A) both reflected that the MPCs coexpressed osteogenesis-, chondrogenesis- and adipogenesis-associated early-stage transcriptional programs at an intermediate level compared with that of primitive-state (cluster 1) and commitment-state (clusters 3, 4, and 5) cells. Transcripts in the osteogenesis program include the upstream transcriptional regulator ID4. Transcripts in the chondrogenesis program include (a) the key TF SCX and (b) the downstream target ECM protein COL11A1. Transcripts in the adipogenesis program include (a) the key TF PPARG and (b) the upstream transactivator CEBPD (Figure 1G). In addition, the predicted TFs in MPCs were related to driving diverse lineage commitment decisions (Figure S2F). This model of MPCs is similar to a model proposed in a previous study, which suggested that multipotent progenitor cells can exist in a lineage-priming state and that lineage-affiliated genes are ‘primed’ for expression later during differentiation [44]. Thus, our results support the concept that lineage priming might occur in MPCs.

**Figure 3.**
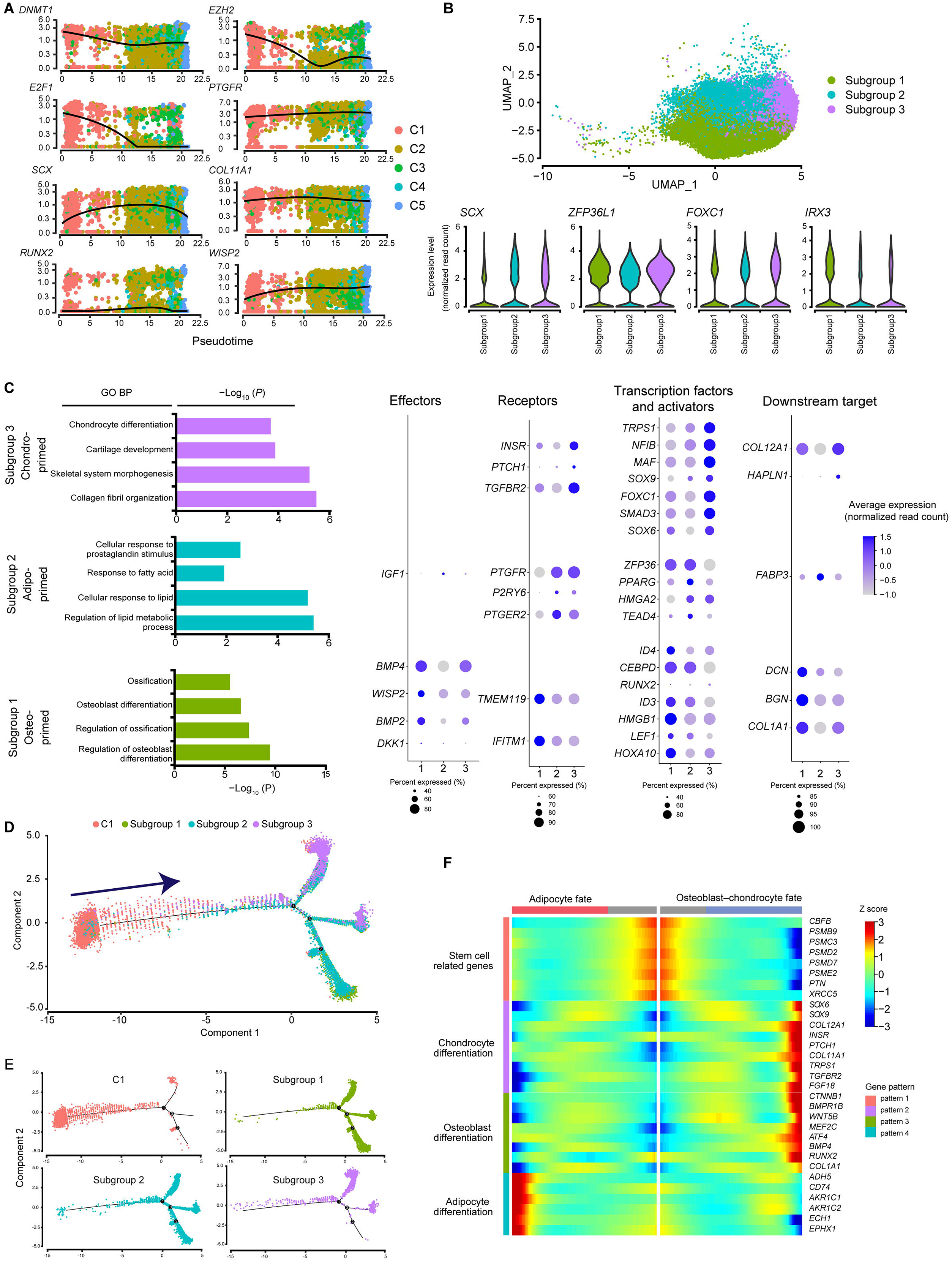
Cluster 2 MPCs are primed towards trilineage orientations. **A.** Relative expression patterns across pseudotime of representative genes for self-renewal maintenance (*DNMT1, EZH2,* and *E2F1*), adipogenesis (*PTGFR*), chondrogenesis (*SCX* and *COL11A1*) and osteogenesis (*RUNX2* and *WISP2*). The dots are colored by cluster name. **B.** Cell type identification on the UMAP plot of MPCs (n = 30,946 cells) (top). Violin plots showing the relative expression levels of genes involved in mesodermal development (*ZFP36L1*, *FOXC1*, *IRX3*, and *SCX*) in each subgroup of MPCs (bottom). **C.** Representative GO terms and corresponding DEGs. Left: Representative GO BP terms enriched with upregulated genes (average log2FC > 0.25) in each subgroup of MPCs. Right: Dot plot showing the relative expression levels of representative genes (from left to right: effectors, receptors, TFs and activators and downstream targets) involved in the indicated terms are shown on a dot plot. The dot size indicates the percentage of cells in the cluster expressing a gene; the shading indicates the relative level of expression (low to high, shown as light to dark). **D.** Developmental trajectory between Up: Pseudotime map of Cluster 1 and each subgroup from MPCs generated by Monocle. Down: Monocle2 plot colorred by each cell cluster identity. **E.** BEAM analysis by Monocle2 indicated different expression patterns during the development of stem-like MSCs to an osteo-chondrogenic or adipogenic fate. MPCs, mesenchymal progenitor cells; BP, biological process; BEAM, branched expression analysis modeling.

Although MPCs are multipotent and possess trilineage differentiation potential, whether these abilities are executed by a single cell population or distinct subgroups of cells is unclear. To explore the diverse progenitors for specific lineages, we subclustered the MPCs by unsupervised clustering via UMAP (**Figure 3,** B). Interestingly, except for the uniform expression of genes involved in mesodermal development (*ZFP36L1, FOXC1, IRX3,* and *SCX*) (Figure 3B and Figure S3D), there were three subgroups showing distinct expression patterns. The controllers of transcriptional programs related to osteogenesis, chondrogenesis, and adipogenesis, such as the TFs RUNX2, SCX, and PPARG, respectively, were expressed at relatively high levels in the aforementioned subgroups. In addition, cells in subgroup 1 expressed genes related to osteoblast differentiation (*IFITM1* and *TMEM119*) and osteoblast progenitor cell proliferation (*COL1A1*, *ID3,* and *ID4)* (Figure 3C). Thus, cells in subgroup 1 were referred to as osteo-primed MPCs. Cells in subgroup 2 expressing genes related to fatty acid metabolism and lipid accumulation (*IGF1, PPARG, FABP3, P2RY6, PTGER2,* and *PTGFR*) (Figure 3C) were referred to as adipo-primed MPCs. Cells in subgroup 3 expressed key genes involved in aspects of chondrogenesis, including chondrocyte differentiation, cartilage development, and collagen fibril organization (*COL11A1, COL12A1, MAF, NFIB, TGFBR2, TRPS1, FGF18,* and *INSR*). Thus, subgroup 3 cells were referred to as chondro-primed MPCs. Therefore, three subgroups of MPCs with differentiation bias, with the possibility of leading to distinct differentiation programs, were identified.

To elucidate the early cell development program of MSCs, we performed developmental trajectory analysis with stem-like APCs (cluster 1), osteo-primed MPCs, adipo-primed MPCs and chondro-primed MPCs (Figure 3D). Two major routes of differentiation from the initial stem-like APCs to the three subgroups of MPCs were revealed, and each route was associated with more than one subgroup of MPCs (Figure 3E). We hypothesized that MPCs in lineage-priming states may adopt stochastic and reversible fates rather than stable states. Then, to investigate the different regulatory patterns of gene expression during this early transition, we performed branched expression analysis modeling (BEAM) on the first bifurcation point with Monocle 2. Hierarchical clustering was performed with the significantly differentially expressed genes during specification, resulting in the identification of four different gene expression patterns during trilineage development (Figure 3F). The genes enriched in pattern 1 were related to stem cell differentiation and were highly expressed in prebranched APCs. These genes included stemness-associated molecules (*CBFB* and *PTN*) and proteasome complex subunits (*PSMB9, PSMC3, PSMD2, PSMD7,* and *PSME2*), which play pivotal roles in the regulation of self-renewal, pluripotency and differentiation of stem cells [45, 46](Figure 3F). Patterns 2 and 3, containing the genes that were upregulated in osteo-chondro-committed precursors, were enriched with chondro-specific genes such as *SOX6, SOX9,* and *COL12A1* [47], as well as the osteo-specific genes *ATF4* and *RUNX2* (Figure 3F) [30, 48, 49]. Pattern 4 contained the genes that were upregulated in adipo-committed precursors (Figure 3F), such as aldo-keto reductases (*AKR1C1* and *AKR1C2*) and *EPHX1*, which are vital for adipocyte differentiation [50, 51]. These results are consistent with the balance towards adipogenesis in favor of osteo-chondrogenesis during MSC commitment [47, 52]. Considering these results collectively, we found that MPCs reasonably exist as a transition state between the waning stem cell program and ongoing progression to osteoblast/chondrocyte/adipocyte fate.

### Prechondrocytes (Cluster 5) specifically harbor immunoregulatory capacity

Genes involved in the chondrogenesis process, including the chondroitin sulfate catabolic process, endochondral bone morphogenesis and skeletal system development, were highly expressed in cluster 5 (**Figure 4,** A). In addition, cluster 5 enriched functional processes involved in the presentation of proinflammatory features, including immunogenicity, complement system activation or inhibition, and myeloid leukocyte activation, as well as anti-inflammatory features, such as suppression of immune cell (e.g., T cell, B cell, NK cell, and DC) proliferation, differentiation and activation (Figure 4, B and C). The predicted TFs, such as IRF1, NFATC2, and NFKB2, and their enriched coexpressed gene sets (Figure 4, D and Figure S4A), are important regulators of the innate and acquired immune responses [53, 54]. These results suggested that cluster 5 cells, referred to as prechondrocytes, possessed immunoregulatory potential.

**Figure 4.**
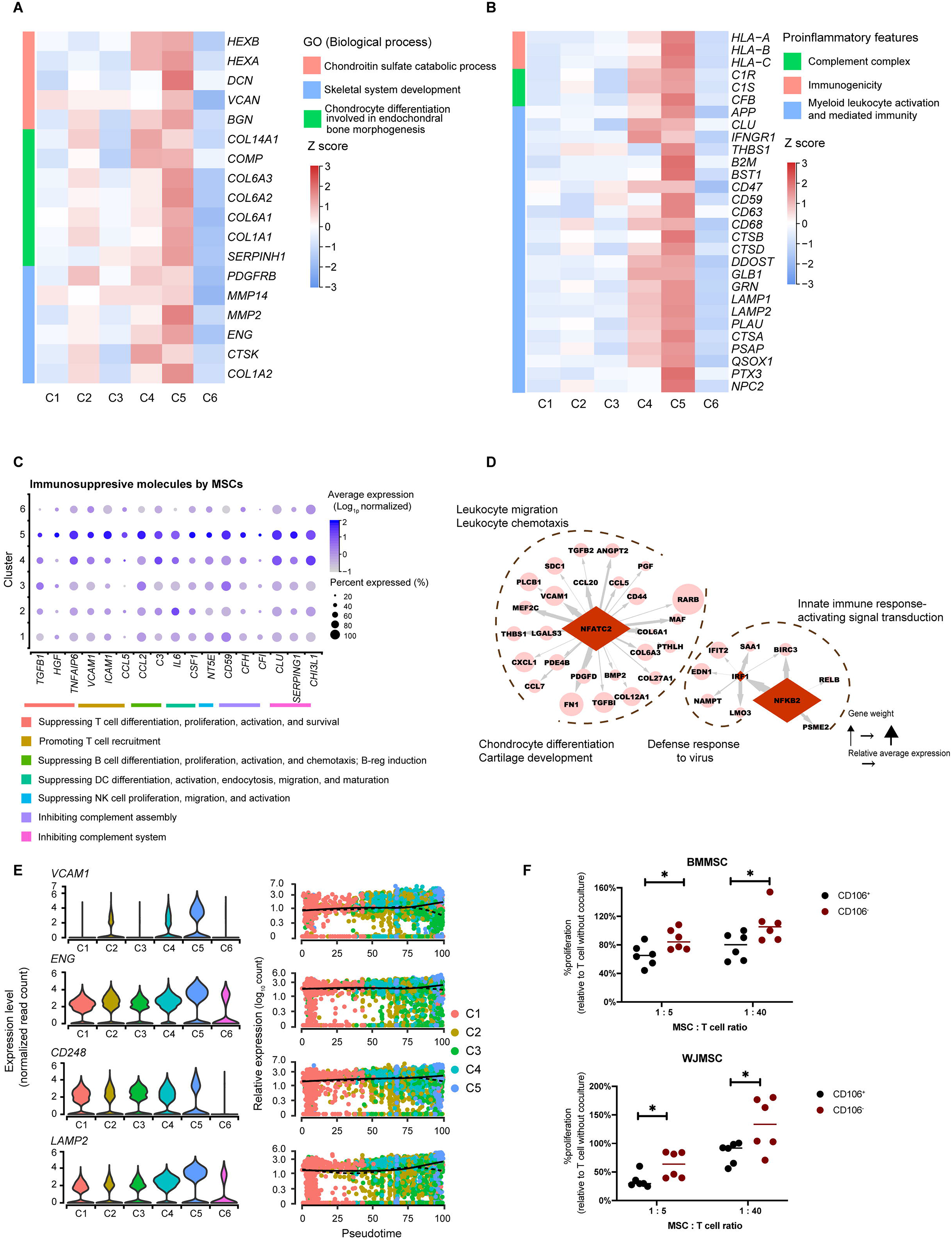
BMMSC-dominant prechondrocytes (Cluster 5) specifically harbor immunoregulatory capacity. **A.** Heatmap showing the relative expression levels of selected genes involved in chondrogenesis in each subpopulation. The color indicates the scaled expression level. **B.** Heatmap showing the relative expression levels of selected proinflammatory genes in each subpopulation. The color indicates the scaled expression level. **C.** Dot plot showing the relative expression levels of selected genes involved in immunosuppression in each subpopulation. The dot size indicates the percentage of cells in the cluster expressing a gene; the shading indicates the relative level of expression (low to high, shown as light to dark). **D.** The potential activated TFs in cluster 5 predicted by SCENIC are shown as diamonds, and their potential downstream targets are shown as circles. **E.** Violin plots (left) and pseudotime trajectories (right) showing the relative expression levels of representative potential markers (CD106/*VCAM1*, *CD248*, CD105/*ENG*, and *LAMP2*). **F.** Proliferation of activated T cells (stimulated with anti-CD2/CD3/CD28 coated microbeads) when co-cultured with CD106^+^ or CD106^−^ MSCs at 1:5 or 1:40. “%proliferative” was measured as the percentage of FSC^high^ Dye^low^ cells in the living cells, and was relative to the positive control group (activated T cells without coculture with MSCs). (n = 6, the data are represented as means ± SDs; \**P* < .05; analyzed by t-test).

In addition to their immunomodulatory profile, cluster 5 cells expressed high levels of genes related to protein processing in the endoplasmic reticulum (*SAR1A, SAR1B,* and *TMEM30A*), protein folding (*HSPA13, DNAJB4,* and *SIL1*), posttranslational protein modification (*PRKCSH, PRSS23, WSB1,* and *RCN1*) and regulation of exocytosis (*LGALS3BP* and *ISLR*) (Figure S4B), consistent with the results of previous studies suggesting that factors in the MSC secretome might perform the major tasks in MSC-mediated immunoregulation [4]. To further clarify how cluster 5 cells regulate inflammation and immune responses, G:Profiler was used to perform cellular component enrichment analysis [55], which showed that most immunomodulatory genes were localized in the extracellular space or extracellular vesicles (Figure S4C). Evaluation of known pathway expression patterns in cluster 5 via gene set variation analysis (GSVA) revealed strong enrichment of pathways such as positive regulation of receptor mediated endocytosis, regulation of endocrine process, organelle membrane fusion and endocrine hormone secretion (Figure S4D). Together, these results suggested that the immunoregulatory effect of cluster 5 MSCs is likely due to their production of exosomes or secretion of soluble factors.

MSCs can modulate the response of immune cells via interaction with lymphocytes, especially T cells, via both the innate and adaptive immune systems to produce anti-inflammatory effects after homing to sites of inflammation *in vivo*. To test the ability of cluster 5 cells to counteract inflammation, we purified cluster 5 MSCs by the surface marker CD106, which was identified as the most significant and specific marker in our data (Figure 4E and S4E). When cocultured with activated CD3^+^ T cells, both CD106^+^ WJMSCs and BMMSCs reduced CD3^+^ T cell proliferation more significantly than did the corresponding CD106^−^ cells (Figure 4F and S5, A to C), indicating that CD106^+^ cluster 5 MSCs exhibited greater anti-inflammatory capacity than other MSC subpopulations. Taken together, these observations suggest that cluster 5 MSCs, identified as prechondrocytes, possessed both proinflammatory and anti-inflammatory potential mediated in a paracrine manner by a variety of secreted soluble factors.

### Cultured MSCs show different properties from primary MSCs by single-cell analysis

To explore the similarity and differences between cultured MSCs and uncultured primary MSCs, we further performed integrated analyses of our data with published scRNA-seq data derived from human primary umbilical cord MSCs (UCMSCs) [12] and BMMSCs [56] as well as cultured endometrial MSCs [57]. By unsupervised clustering, six similar clusters were observed in cultured endometrial MSCs, BMMSCs, and WJMSCs (Figure S6A), including stem-like APCs (cluster 1), MPCs (cluster 2), preadipocytes (cluster 3), pre-COs (cluster 4), prechondrocytes (cluster 5), and SMCs (cluster 6). This result suggested that cultured MSCs from different tissues shared similar subpopulation compositions. Interestingly, UCMSCs were mainly composed of cluster 2 cells, while primary BMMSCs were mainly composed of cluster 2 cells as well as cells from a novel BMMSC specific cluster (cluster 7) (Figure S6A). This primary BMMSC specific cluster (cluster 7) highly expressed HSC-niche factor genes, including *CXCL12, VCAM1, and ANGPT1*, (Figure S6B), and was thus referred to as the “HSC-niche support cluster”. The discrepancy in cluster groups between cultured and uncultured MSCs suggested that primary BMMSCs may lose their original gene expression activity related to HSC-niche support after *ex vivo* culturing, which is consistent with a previous finding that MSC lost their HSC-niche function during culture [58]. In addition, the characteristic surface marker for MSCs, including CD73*/NT5E,* CD90*/THY1,* and *CD44*, were expressed higher in cultured cells than primary MSCs both at the expression level and in terms of the percentage of positive cells (Figure S6C). This suggested that these MSC characteristics might increase during culture. Taken together, our findings not only uncover the molecular and functional heterogeneity of cultured MSCs, but also pave a way for exploring the distinct heterogeneous characteristics between cultured and uncultured MSCs.

## Discussion

MSCs are considered promising candidates for cell-based regenerative medicine due to their self-renewal capacity, multilineage differentiation potential, paracrine effects, and immunosuppressive properties [59, 60]. However, whether and to what extent the MSC population contains heterogeneous subpopulations associated with diverse functions and characteristics remains largely unknown. In this study, we performed high-throughput scRNA-seq and a comprehensive analysis on *ex vivo*-expanded human BMMSCs and WJMSCs, which represent cell sources from adult and neonatal tissues, respectively. Our study identified the inherent cellular composition of MSCs, which included a stem-like active proliferative cell subpopulation, a multipotent progenitor subpopulation, specific adipocyte and osteo-chondrocyte precursor subpopulations, and an immunoregulatory prechondrocytes subpopulation, as additionally providing a reconstruction of the transcriptional hierarchies of these subpopulations.

MSCs exhibit stemness characteristics *in vitro*, expanding rapidly and maintaining their morphology for up to 10 passages. However, there is still a lack of evidence to identify the stem cell populations among MSCs. In our data, cluster 1 cells specifically expressed *CSPG4/MCAM/NES* and stemness signatures and exhibited negative regulation of differentiation pathways. Combining these properties with its location at the apex of the developmental trajectory, cluster 1 was referred to as the stem-like cell cluster. Interestingly, MSCs were reported to be derived *in vivo* from NESTIN^bright^ NG2/CSPG4^+^ periarteriolar mesoderm progenitor cells, which possess the stemness characteristics of self-renewal through serial transplantations and multilineage mesodermal differentiation potential [21–23]. This suggests that stemness might be maintained in long-term culture as the cluster 1 subpopulation. Moreover, stem cells in long-term culture show a strong proliferative ability to support successive rounds of replication and passaging without differentiation [61, 62], consistent with the highly proliferative phenotype of cluster 1 cells. In addition, while cluster 1 was closely related to NG2^+^ periarteriolar cells, cluster 1 (APCs), 3 (preadipocytes), and 5 (prechondrocytes) have a comparatively higher similarity with LEPR^+^ perisinusoidal cells (Figure 2C). As LEPR^+^ cells serve as the major source of cartilage and adipocytes in adult mouse bone marrow [63], the comparative similarity is supportive of the adipogenesis or chondrogenesis potentials of these three clusters (cluster 1, 3, and 5).

Lineage priming is a molecular model of stem/progenitor cell (S/PC) differentiation in which S/PCs express low levels of a subset of genes associated with the differentiation pathways to which they can commit. Thus, they are “primed” for expression later during differentiation [64]. This concept has been widely used to explain the stochastic differentiation ability of hematopoietic stem cells. A similar process might occur with MSCs. Previous studies have shown that MSCs simultaneously express markers of more than one mesenchymal lineage [46, 64], suggesting the existence of a lineage-priming state in MSCs. However, the transcriptome pattern of lineage priming in MSCs and the relationships between lineage priming and lineage specification are incompletely understood. With the support of advanced scRNA-seq techniques, we revealed that MPCs, a subpopulation of MSCs, coexpressed lineage-associated TFs, markers and receptors, suggesting the presence of a lineage-priming state specifically in MPCs. Furthermore, subclustering identified three lineage-biased subgroups, suggesting that lineage specification began in multipotent progenitors with lineage priming. Together, our studies extended the concept of lineage priming to MSCs and shed light on the potential developmental continuum connecting stem cells to downstream precursors.

MSCs exhibit proinflammatory and anti-inflammatory properties [4, 65, 66]. However, whether these properties were exerted by homogeneous MSCs or by a distinct subset of cells remains elusive. Here, we found that the immunomodulatory function of MSCs was likely executed by a specific subpopulation (cluster 5) instead of the entire MSC population. This subpopulation of cells specifically expressed genes related to proinflammatory and anti-inflammatory signatures, supporting the immunoregulatory plasticity of MSCs. We further identified specific surface markers for this cluster, including CD106*/VCAM1, CD47, CD248,* CD87*/PLAUR,* etc. CD106^+^ MSCs derived from placental chorionic villi were demonstrated to be more effective in modulating T helper subsets [67, 68]. However, it is still unknown whether the immunoregulatory signatures are limited to CD106^+^ cells. Our study showed the specificity of this immunoregulatory subpopulation, which was identified as prechondrocytes and located at the end of the differentiation paths. Moreover, previous studies showed that mature chondrocytes can exert an anti-inflammatory effect, for example, primary chondrocyte-derived exosomes can prevent osteoarthritis progression via expression of anti-inflammatory cytokines [69]. This research supports the findings of our study that prechondrocytes can express complex and varied immune signatures and likely produce exosomes or secrete factors that are the basis of the immunomodulatory function of MSCs. A recent study performing scRNA-seq on primary WJMSCs revealed distinct subpopulations defined by enrichment of terms related to proliferation, development, and inflammation response, but the specific markers and functional modulators for subpopulation identification need further investigation. Moreover, increasing evidences have shown that the immunoregulation functions exerted by MSCs were cell-contact dependent and/or produce various immunoregulatory and growth factors [4, 65, 66, 70]. As the specific immunomodulatory subpopulation (cluster 5) was identified in our study, it is important to explore the pattern and mechanisms of its immunoregulatory effect in detail in the future.

MSCs from neonatal tissues show higher proliferative capacity than MSCs from adult tissues, and MSCs derived from bone marrow significantly inhibit allogeneic T cell proliferation [19, 71]. By scRNA-seq analysis, we found that WJMSCs contained a higher percentage of proliferative stem-like cells (cluster 1) compared to BMMSCs (17.3% *vs* 5%), supporting the biological superiority of WJMSCs in expansion [71, 72]. On the other hand, the superiority of BMMSCs in immunomodulation [72, 73] could be related to the dominance of prechondrocytes subpopulations (Figure S2C, BMMSCs, 98.84% *vs* WJMSCs, 1.16%) and the increased proportion of CD106^+^ cells (Figure S4E, BMMSCs, 72.73%±24.66 *versus* WJMSCs, 23.78%±11.29). Therefore, the tissue-specific MSC characteristics can be potentially ascribed to the different proportions of functional clusters.

To our knowledge, scRNA-seq studies of different types of MSCs have been published, including the out-of-thaw MSCs, iPSC-derived MSCs as well as the *in vivo* primary MSCs. The potential phenotypic signatures of MSCs were varying among these studies with some important differences in emphasis. For example, contrasting pre-freeze and out-of-thaw samples, Medrano-Trochez et. al., 2020 found that out-of-thaw MSCs expressed higher levels of cholesterol/steroid biosynthesis and regulation of apoptosis, but lower levels of cytokine signaling, cell proliferation, and cell adhesion [74]. When investigating the gene regulatory networks during chondrogenesis from hiPSC, Wu et.al., 2021 found that inhibiting WNTs and MITF could enhance the yield and homogeneity of hiPSC-derived chondrocytes [75]. A unique adipogenic lineage precursors (MALPs) was identified in primary BM-MSCs that played critical roles in maintaining marrow vasculature and suppressing bone formation as an important part of niche cells [76]. Huang et al. profiled the single-cell transcriptomes of 361 UC-MSCs from 7 samples under different conditions, revealing that hMSCs had limited heterogeneity [9]. However, the cell number of each sample was relatively small (~ 50 cells/sample) and they only analyzed the samples individually. The conclusion that MSCs had limited heterogeneity depended only on the similar 4 subclusters and gene expression patterns identified in each individual sample. Moreover, they also mentioned that the limited heterogeneity in these UC-MSCs was strongly associated with the dominant cell cycle effect on MSCs. Comparatively, our study dissected the heterogeneity by integration analysis of a large number of MSCs and intentionally removed the cell cycle effects, which would be more conducive for subpopulation identification. Additionally, cultured conditions might influence the expansion and differentiation ability of MSCs, which will also impact the heterogeneity of MSCs. Pattappa et al. have reported that MSCs expanded with normoxia (20% oxygen) had more rapid initial proliferation but contained a greater proportion of senescent cells than those under hypoxia (5% or 2% oxygen). These phenomena were associated with the metabolic profiles from glycolysis [77]. Xie et al. also indicated that the metabolic profile of MSCs impactedtheir functional heterogeneity [78]. Thus, it should be noted that the effects of cellular metabolism should be considered during MSC culture and application.

In summary, we performed a comprehensive investigation of the heterogeneity of MSCs and discovered distinct subpopulations with specific characteristics, including stem-like proliferative cells, multipotent progenitors and specific lineage precursors, as well as immunomodulatory prechondrocytes subsets. We constructed the developmental hierarchies of cellular subpopulations among cultured MSCs for the first time. These transcriptional profiles identified MSC subsets and related them to specific markers that could be used to purify functional subpopulations for more specific and effective therapeutic applications.

## Materials and methods

### Isolation and culture of WJMSCs and BMMSCs

For isolation of WJMSCs and BMMSCs, fresh human umbilical cords and bone marrow samples were obtained (Table S1). The umbilical cord was cut down into smaller segments. Then, the arteries and veins were removed and the remaining parts were immersed in a stem cell culture medium. BMMSCs were obtained by bone marrow puncture aspiration of the iliac crest cavity from young children with cerebral palsy. Mononuclear cells were collected by Ficoll-based density gradient centrifugation and cultured in T75 flasks at a density of 160,000/cm^2^ in MEM alpha basic (C12571500BT MEM α, Nucleosides MEM, Invitrogen, Carlsbad, CA) culture medium supplemented with penicillin and streptomycin (Gibco, Carlsbad, CA) and 10% fetal bovine serum (IVGN-10099141, Gibco). Cells were cultured at 37°C in a humidified atmosphere with 5% CO2. Cells were passaged and trypsinized with 0.25% trypsin/EDTA at 80%–90% confluence. MSCs intended for functional assays were harvested between passage 2 and passage 5. Passage 6 or 7 cells were used for subsequent scRNA-seq analysis.

### Osteogenic lineage differentiation and staining analysis

MSCs were subcultured in 6-well plates at an initial density of 2 × 10^4^/cm^2^ with standard expansion medium. When the cells reached 60%–80% confluence, the medium was changed into 2ml human osteogenic differentiation medium (HUXUC-90021, Cyagen Biosciences, China). The medium was refreshed every 3 days. After 2– 4 weeks’ differentiation, cells were rinsed by Dulbecco’s Phosphate-Buffered Saline (DPBS, C14190500CP, Invitrogen) and fixed with 4% paraformaldehyde. Then the cells were stained with 1ml Alizarin Red S solution for 3–5 min. Calcified matrix was stained red with Alizarin Red S, indicating the deposition of calcified matrixes on the osteogenic differentiated hMSCs.

### Adipogenic lineage differentiation and staining analysis

MSCs were subcultured in 6-well plates at an initial density of 2 × 10^4^/cm^2^ with standard expansion medium. When the cells were 100% confluent, the medium was changed into 2ml osteogenic differentiation medium A (HUXUC-90031, Cyagen Biosciences, China). After three days, the medium was changed into 2ml Adipogenic Differentiation Medium B. After alternating the A and B fluids 3–5 times (12–20 days), B-liquid was used for 4–7 days until a sizable amount of large and round lipid droplets emerged. During B-fluid maintenance culture, fresh B-liquid was replaced every 2–3 days. Then, cultured MSCs were rinsed by DPBS 1–2 times and were fixed for 30min at room temperature with 4% paraformaldehyde. After the DPBS rinse, cells were stained with 1ml Oil Red O solution for 30 min. Oil Red O imparts red-orange color to the lipid droplets.

### Chondrogenic lineage differentiation and staining analysis

MSCs for hondrogenic differentiation were cultured with chondrogenic differentiation medium (HUXUC-90041, Cyagen Biosciences, China) for 14 days. The chondrogenic aggregates were fixed with 10% formalin for 20 min and stained with Alcian Blue 8GX for 30 min.

### Flow cytometry

MSC samples were examined by flow cytometry analysis with the anti-human antibodies, including CD73 (CD73-PE, 12-0739-41, eBioscience, Thermo Fisher Scientific, Waltham, MA), CD90 (CD90-PE, 12-0909-42, eBioscience), CD105 (CD105-APC, 17-1057-41, eBioscience), CD34 (CD34-APC, 17-0349-42, eBioscience), CD45 (CD45-PE, 12-0459-42, eBioscience), CD11b (CD11b-APC, 101212, BioLegend, San Diego, CA), CD19 (CD19-FITC, 555412, BD, Franklin Lake, NJ) and HLA-DR (HLA-DR-FITC, 11-9956-41, eBioscience). Cells were harvested and re-suspended in a staining buffer (2% FBS in DPBS), and were subsequently incubated with corresponding antibodies at 4℃ for 30 min avoiding light. After the samples were washed with DPBS and re-suspended in staining buffer, they were run on a Moflo XDP (Bechman). For each sample, more than 8000 events were acquired.

### T cell proliferation assay

Briefly, T cells were purified from peripheral blood samples by the negative selection EasySep^TM^ Human T cell Enrichment Kit (Catalog #17951, StemCell Technologies, Vancouver, Canada). Enriched T cells were stained with Cell Proliferation Dye eFluor^TM^ 450 (Catalog #65-0842, eBioscience) to assess cell proliferation. Dye eFluor^TM^ 450-labeled T cells (2.5 × 10^4^) were stimulated with anti-CD2/CD3/CD28 coated microbeads (Pan T Cell Activation Kit; Miltenyi Biotech, Bergisch Gladbach, Germany) or uncoated microbeads as a negative control in a 1:10 bead:T cell ratio. These cells were co-cultured with allogeneic CD106^+^ or CD106^−^ MSCs, which had been previously seeded in 96-well plates (5000 or 625 cells/well). The percentage of T cell proliferation was measured after 3.5 days in a Moflo XDP (Bechman) and calculated by the percentage of FSC^high^Dye^low^ cells in the living cells gated by FSC/SSC. More than 8000 events were acquired for analysis. The data was normalized with respect to the percentage of activated T cells without coculture with MSCs.

### ScRNA-seq library preparation and sequencing

Validated WJMSCs and BMMSCs were collected and re-suspended at 1 × 10^6^ /ml in DPBS with 0.04% BSA. Cells with higher aggregation rate (measured by a Countstar cell count and analysis system) were filtered to remove the cell aggregates. The cell suspensions (> 90% living cells examined by Countstar) were loaded on a Chromium Single Cell Controller (10 × Genomics). The scRNA-seq libraries were sequenced on Illumina Hiseq X-ten platform with a 150-bp paired-end read length.

### ScRNA-seq data processing

Raw sequencing data were processed by the Cell Ranger 3.0.2 pipeline with default parameters. Each sample was aligned to GRCH38 by the “Cell Ranger Count” function to get the raw gene expression matrices. These matrices were further analyzed by Seurat (v3.0.2) for quality control and downstream analysis [79]. Low-quality cells that had less than 200 genes per cell and less than 3 cells per gene were discarded. Then to remove the outliers, cells were kept based on stringent criteria: 1000 < genes per cell < 6500, and percentage of mitochondrial genes < 0.05. After quality control, a total of 61,296 cells were retained.

### Dimension reduction, clustering and identification of differentially expressed genes

The top 4000 most highly variable genes from each sample were selected for data integration. In our scRNA-seq experiments, six MSC samples were sequenced in three batches. CCA was applied to remove the batch effect for data integration. Next, the top 20 PCs of the integrated data were selected for PCA, UMAP analysis and graph-based clustering (resolution = 0.15) to identify distinct subpopulations. DEGs were identified by the ‘FindAllMarkers’ function in Seurat (min.pct = 0.25, thresh.use = 0.25). Metascape [80] were used for gene ontology (GO) and pathway enrichment analysis.

### Pseudotime analysis

The Monocle2 package (version 2.8.0) [81] were used to determine the pseudotime developmental relationships of each cluster in MSCs. We used top 3000 HVGs identified by Seurat to sort cells in pseudo-time order. The ‘DDRTree’ function was applied to reduce dimensions to infer the potential developmental path, and the ‘differentialGeneTest’ function was applied to identify differential expression genes along pseudo-time order. The remaining parameters were default.

### RNA velocity

RNA velocity is introduced to calculate the spliced and unspliced RNAs to indicate the transcriptional kinetic activity. A loom file with counts divided in to spliced / unspliced / ambiguous of each gene in each cell was generated by velocyto.py on the BAM file from the CellRanger analysis. Only cells identical to the Seurat object (cluster1– cluster5) were retained for downstream analysis. Then RNA velocity was estimated by velocyto.R with default settings. The velocity fields were projected on to the UMAP embedding from the Seurat analysis.

### Integrated analysis of scRNA-seq datasets

Seurat (v3.0.2) was applied to integrate the public scRNA-seq datasets with our data. The top 4000 featured genes that were repeatedly variable across datasets were used to identify anchors across batches with the ‘FindIntegrationAnchors()‘ function. The anchors were used to guide integration across multiple datasets with the‘IntegrateData()‘ function. The corrected data (integrated assay) was used for downstream analysis.We regressed out the difference between the G2/M and S phase scores with the ‘ScaleData’ function to remove cell cycle effects.

### TF regulon analysis

TF regulons were analyzed using Single Cell Regulatory Network Inference and Clustering (SCENIC v1.1.2-2) workflow in R [82]. Normalized data from Seurat were used to generate the regulon activity score of TFs by default parameters. The average activity level of regulons in each cluster were calculated to show the main regulatory changes of different clusters through hierarchical clustering. Regulons showing the significant difference in average activity between clusters were selected shown by heatmap.

### Data comparison

The publicly available datasets of LEPR+ and NG2+ cells was downloaded from GEO database with accession number GSE128423 (https://www.ncbi.nlm.nih.gov/geo/query/acc.cgi?acc=GSE128423). The publicly available datasets of DLL1+ PXM, LTM and ESM can be found at SRA under BioProject PRJNA319573 (https://www.ncbi.nlm.nih.gov/bioproject/PRJNA319573/). The publicly available datasets of osteoblasts and chondrocytes was downloaded from GEO database with accession number GSE106292 (https://www.ncbi.nlm.nih.gov/geo/query/acc.cgi?acc=GSE106292). The publicly available datasets of BMMSCs was downloaded from GEO database with accession number GSE147287 (https://www.ncbi.nlm.nih.gov/geo/query/acc.cgi?acc=GSE147287). The publicly available datasets of cultured endometrial MSCs was downloaded from GEO database with accession number GSE149651 (https://www.ncbi.nlm.nih.gov/geo/query/acc.cgi?acc=GSE149651). The publicly available datasets of primary umbilical cord MSCs can be found at SRA under BioProject PRJNA643879 (https://www.ncbi.nlm.nih.gov/bioproject/?term=PRJNA643879). The same single-cell analysis approach in this study was applied to public scRNA-seq data. Then we quantified the correlation of single-cell clusters based on average gene expression of the typical gene signatures related to specific characteristics. Moreover, we performed Combat to remove the batch effects [83].

### Statistical analysis

Statistical analysis of the data was performed by a one-way analysis of variance (ANOVA) followed by Tukey’s post-hoc test among those with more than two groups. Statistical analysis of the data was performed by t test between two groups. *P* values < 0.05 were considered statistically significant. Analyses were performed using R packages or GraphPad Prism 8.

## Ethical statement

This study was approved by the ethics committee of 3rd medical center, General Hospital of Chinese PLA. All subjects provided written informed consent according to the Institutional Guidelines.

## Data and materials availability

Raw data from RNA-seq analysis have been deposited in the Genome Sequence Archive (GSA) [84] under accession numbers: HRA000220 (https://ngdc.cncb.ac.cn/gsa-human/s/LJ7vdOsP). The expression matrix reported in this paper has been deposited in the OMIX, China National Center for Bioinformation / Beijing Institute of Genomics, Chinese Academy of Sciences (https://ngdc.cncb.ac.cn/omix: accession no.OMIX745) [85]. All other data supporting the findings of this study are available from the corresponding authors on reasonable request.

## CrediT author statement

**Chen Zhang:** Investigation, Data curation, Formal analysis, Validation, Visualization, Writing - original draft, Writing - review & editing. **Xueshuai Han:** Investigation, Data curation, Formal analysis, Visualization, Writing - original draft, Writing - review & editing. **Jingkun Liu:** Validation, Investigation. **Lei Chen:** Validation. **Ying Lei:** Validation. **Kunying Chen:** Validation. **Jia Si:** Formal analysis. **Tian-yi Wang:** Writing - review & editing. **Hui Zhou:** Resources. **Xiaoyun Zhao:** Resources. **Xiaohui Zhang:** Resources. **Yihua An:** Resources. **Yueying Li:** Conceptualization, Writing - original draft, Writing - review & editing, Project administration, Funding acquisition. **Qian-Fei Wang:** Conceptualization, Writing - review & editing, Supervision, Funding acquisition. All authors read and approved the final manuscript.

## Competing interests

The authors have declared no competing interests.

## Acknowledgments

We would like to thank Yutian Deng (Beijing Institute of Genomics, Chinese Academy of Sciences (CAS)) and Ting Li (Institute of Genetics and Developmental Biology, CAS) for flow cytometry sorting and analysis. We thank Xing Zhu (Qilu hospital of Shandong University (Qingdao)) for chondrocyte embedding and paraffin sectioning. This work was supported by the National Natural Science Foundation of China (Grant No. 81890992 to Qian-fei Wang, No. 81770109 and 81970108 to Yueying Li), Youth Innovation Promotion Association of Chinese Academy of Sciences (Grant No. 2017142 to Yueying Li).

