## Supplemental legend for "Single-cell Transcriptomic Analysis Reveals the Cellular Heterogeneity of Mesenchymal Stem Cells"

### Supplementary material

#### Figure S1 Characteristics of MSCs and quality control of scRNA-seq data

**A.** A representative flow diagram of the expression of positive markers (CD73, CD90, and CD105) and negative markers (CD34, CD45, CD11b, CD19 and HLA-DR) in WJMSCs. **B.** A representative flow diagram of the expression of positive markers (CD73, CD90 and CD105) and negative markers (CD34, CD45, CD11b, CD19 and HLA-DR) in BMMSCs. **C.** Histogram showing the proliferation ability as evaluated by the ratios of the cell counts on day 3 and day 5 relative to that on day 0. (n = 2, the data are expressed as the mean ± SD. *** $P < .005$; analyzed by t-test.) **D.** Histogram showing the average diameters as analyzed with a Countstar instrument. (n = 6, the data are expressed as the mean ± SD. * $P < .05$; analyzed by t-test.). **E-G.** Box plot showing the number of UMI counts (E), expressed genes (F) and the percentage of mitochondrial genes (G) from single-cell transcriptomic data in WJMSCs and BMMSCs. **H.** Histogram showing the expression patterns of positive marker genes (CD73/*NT5E,* CD90/*THY1*, CD105/*ENG* and *CD44*). The data are expressed as the mean ± SD. UMI, unique molecular identifier; scRNA-seq, single cell RNA-sequencing.

#### Figure S2 Cellular taxonomy of MSCs by scRNA-seq.

**A.** Pie chart showing the cell number and proportion of each subpopulation. **B.** Sample distribution. Left: UMAP plot showing the cell distribution of the two MSC sources (WJMSCs and BMMSCs). Right: Histogram showing the proportion of each subpopulation from the two MSC sources (WJMSCs and BMMSCs). **C.** Histogram showing the percentage of the two MSC sources in each subpopulation. **D.** Dot plot showing the relative expression levels of positive marker genes (CD73/*NT5E,* CD90/*THY1*, CD105/*ENG*, and *CD44*). The dot size indicates the proportion of cells in the cluster expressing a gene; the shading indicates the relative level of expression (low to high, shown as light to dark). **E.** UMAP plot showing the cell distribution of the six individual samples. **F.** Heatmap showing activated TFs predicted by SCENIC. **G.** Heatmap of the correlation matrix of transcriptome between our MSC subpopulations and specific cells from published dataset [34], including osteoblasts and chondrocytes. **H.** Violin plots showing the expression levels of selected genes (*OMD*, *ASPN*, *GPM6B*, *IFITM1, GPNMB, COL6A3, COL6A1,* and *ECM1*) in each cluster. **I.** Monocle2 plot colored by each cell cluster identity. **J.** Velocity field projected onto the UMAP embedding. Arrows show the local average velocity evaluated on a regular grid. TFs, transcription factors.

**Figure S3 Cluster 1 cells exhibited a stem-like transcriptional signature.**

**A.** Heatmap of the correlation matrix between the transcriptome of each subpopulation and cells from published datasets [43], including DLL1$^{+}$ PXM, LTM and ESM. **B.** Dot plot showing the relative expression levels of the activated transcription factors (*CTCF, EZH2, E2F8, PBX3, MYBL2, E2F1, and TFDP1*) in cluster 1 predicted by SCENIC. **C.** Violin plots showing the expression levels of stemness-associated markers (*CD9, CD44, ITGB1, SDC4*, and *ITGAV*) in each cluster. **D.** Violin plots showing the expression levels of genes involved in mesodermal development (*SCX, ZFP36L1, FOXC1,* and *IRX3*) in each cluster.

**Figure S4 Cluster 5 cells exhibited an immunoplasticity transcriptional signature.**

**A.** Dot plot showing the relative expression levels of predicted TFs in cluster 5 (*NFATC2, NFKB2,* and *IRF1*). The dot size indicates the proportion of cells in the cluster expressing a gene; the shading indicates the relative level of expression (low to high, shown as light to dark). **B.** Dot plot showing the relative expression levels of selected genes involved in the indicated biological processes in cluster 5. The dot size indicates the proportion of cells in the cluster expressing a gene; the shading indicates the relative level of expression (low to high, shown as light to dark). **C.** GO analysis with g:Profiler showing CC terms enriched with differentially expressed genes in cluster 5. The gene count is indicated by the dot size, and the gene ratio is indicated by the color of the dot (blue, low ratio; red, high ratio). **D.** Histogram showing representative enriched GO terms involved in protein processing in cluster 5 identified by GSVA. **E.** Representative flow cytometry plots and statistical data of CD106/*VCAM1* expression levels in MSCs from WJMSCs and BMMSCs (n = 7, the data are expressed as the mean ± SD. *** $P < .005$; analyzed by t-test.). CC, cellular component; GSVA, gene set variation analysis.

**Figure S5 The immunoregulatory capacity of cluster 5 was supported by the stronger inhibition on activated CD3$^{+}$ T cells.**

**A.** A representative flow diagram (up) and histogram (bottom) showing T cell proliferation with and without CD2/CD3/CD28 stimulation (n = 8, the data are expressed as the mean ± SD. *** $P < .005$; analyzed by t-test; normalized with respect to the percentage of non-activated T cells). **B.** A representative flow diagram of T cell proliferation co-cultured with $CD106^+$/$CD106^-$/non-sorted MSCs with and without activation. **C.** Histogram depicting T cells proliferation stimulated with anti-CD2/CD3/CD28 coated microbeads co-culturing with $CD106^+$/$CD106^-$/non-sorted MSCs at 1:5 or 1:40, the data were analyzed by dye loss and FSC/SSC gating, respectively. The data were relative to the positive control group, that was the percentage of activated T cells without coculture with MSCs (n = 6, the data are expressed as the mean ± SD. *** $P < .005$; analyzed by one-way ANOVA followed by Tukey's post-hoc test).

**Figure S6 The integration analysis between our dataset and published dataset.**

**A.** UMAP plot showing the source distribution from the integration analysis, including the BMMSCs and WJMSCs from our data and the cultured human endometrial MSCs [57] , uncultured human UC-MSCs [12], and primary BMMSC [56] from published scRNA-seq data. **B.** Violin plots showing the expression levels of genes involved in mesodermal development (*LEPR, CXCL12,* and *ANGPT1*) in each cluster. **C.** The feature plot shows the expression of MSC markers (CD73/*NT5E,* CD90/*THY1,* and *CD44*) in each source.

**Table S1 Sample information.**

**Table S2 Single cell metadata table**
