## Supplementary figures and images for "Single-cell Transcriptomic Analysis Reveals the Cellular Heterogeneity of Mesenchymal Stem Cells"

### Supplemental Figure 1

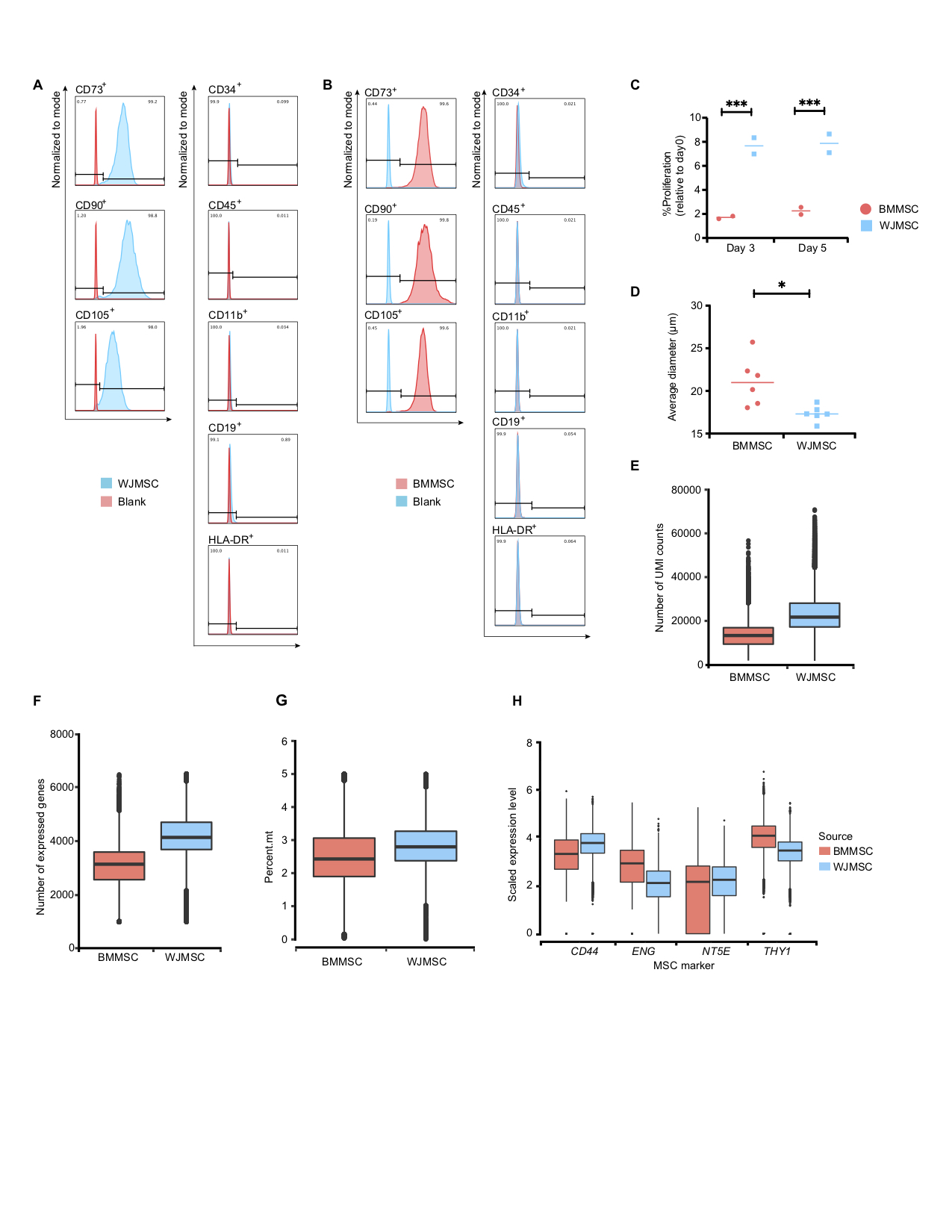

### Supplemental Figure 2

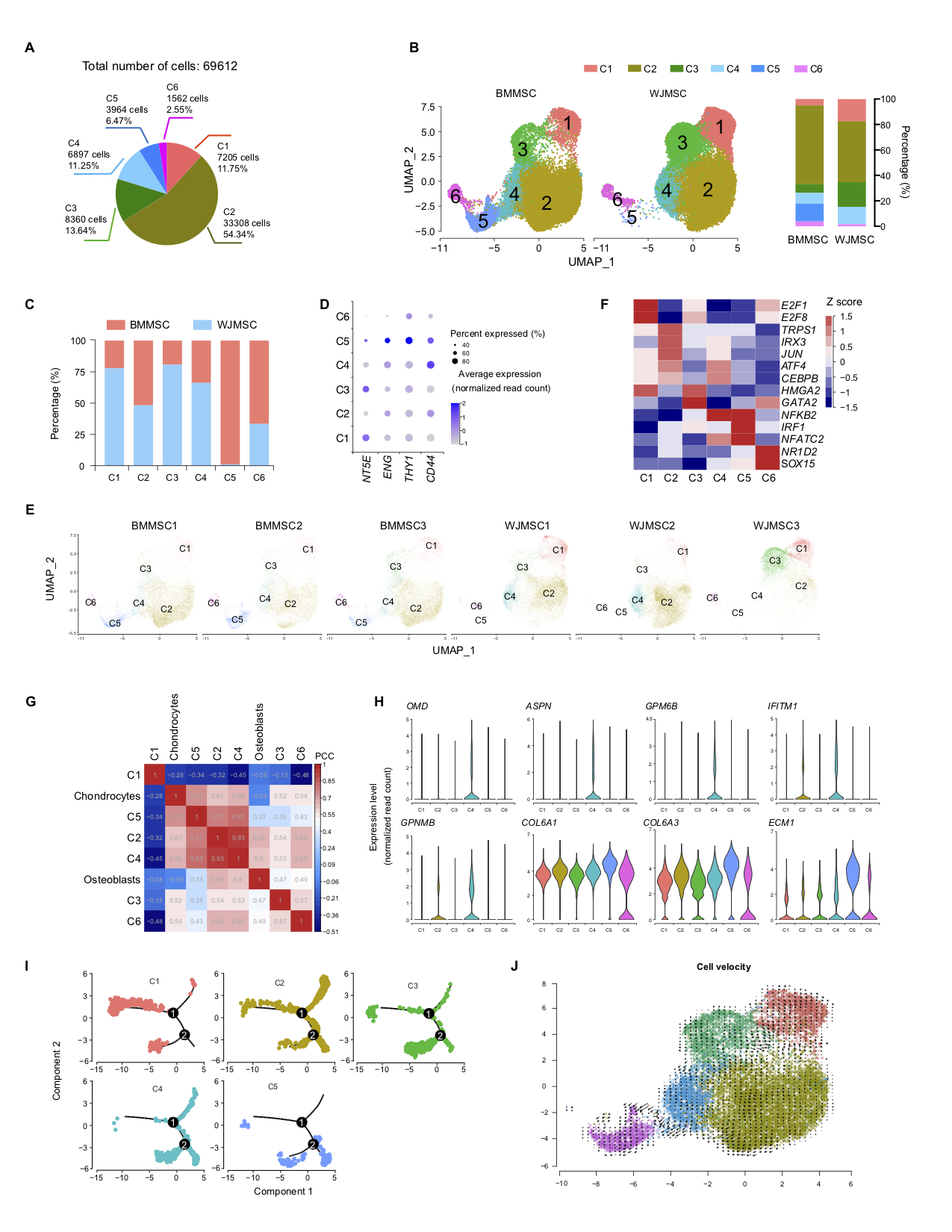

### Supplemental Figure 3

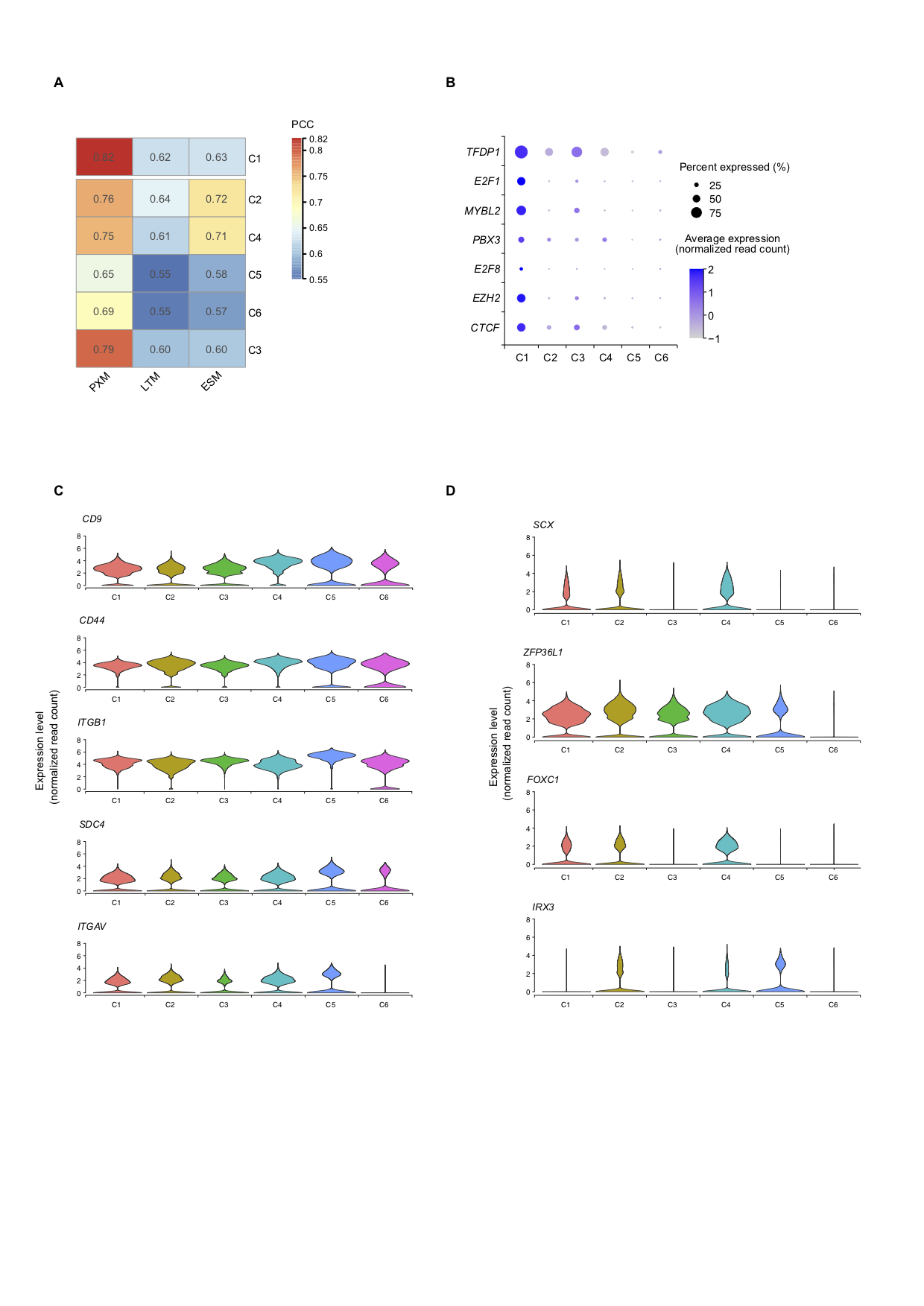

### Supplemental Figure 4

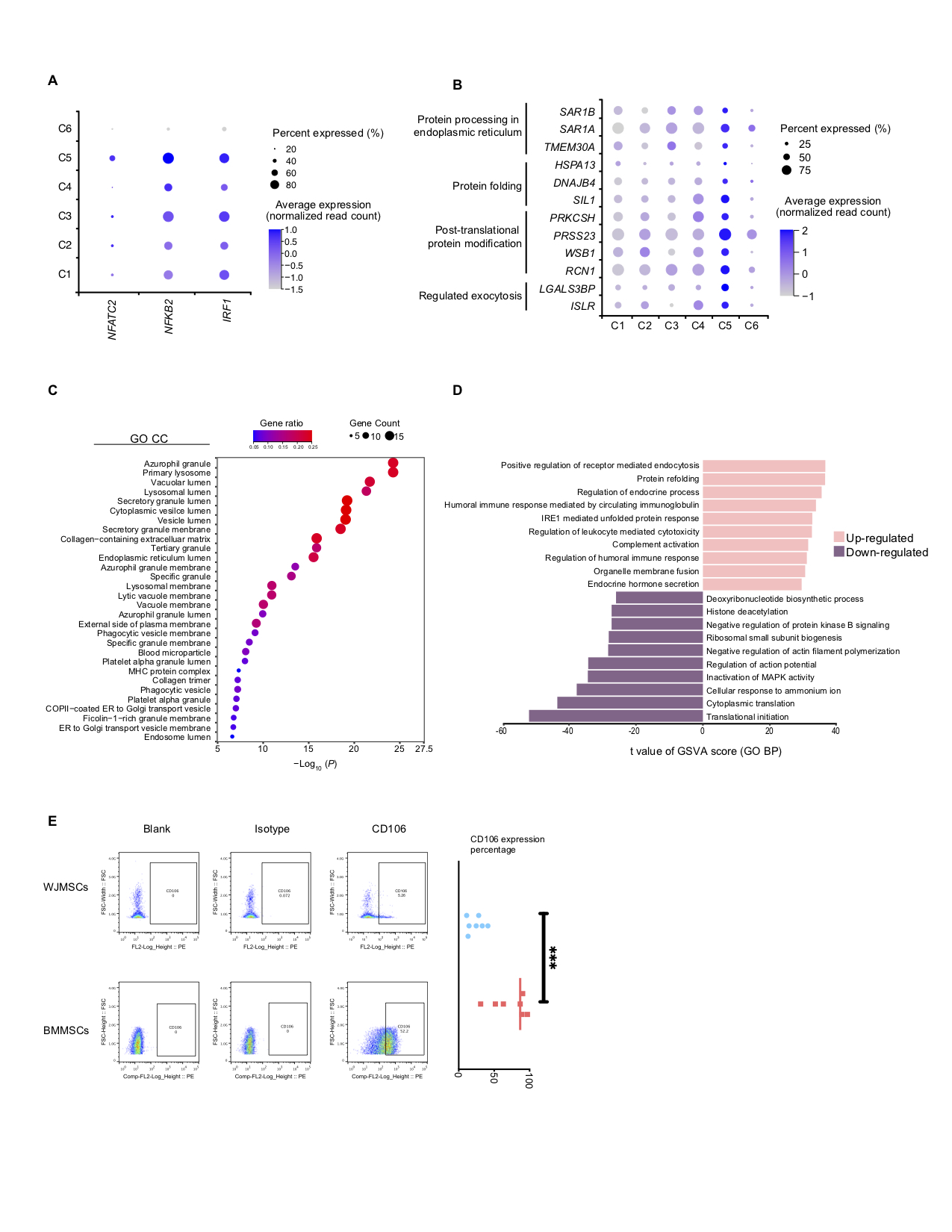

### Supplemental Figure 5

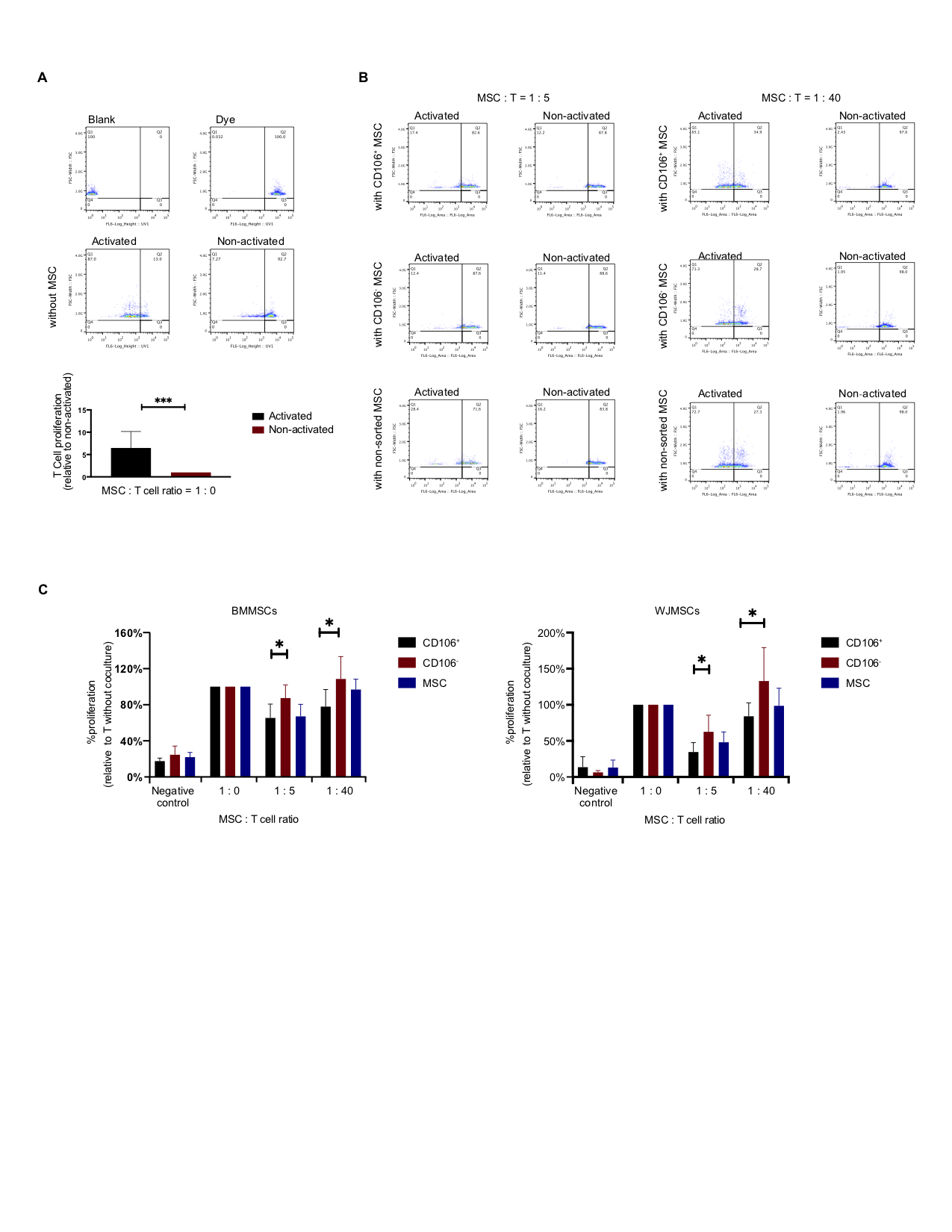

### Supplemental Figure 6

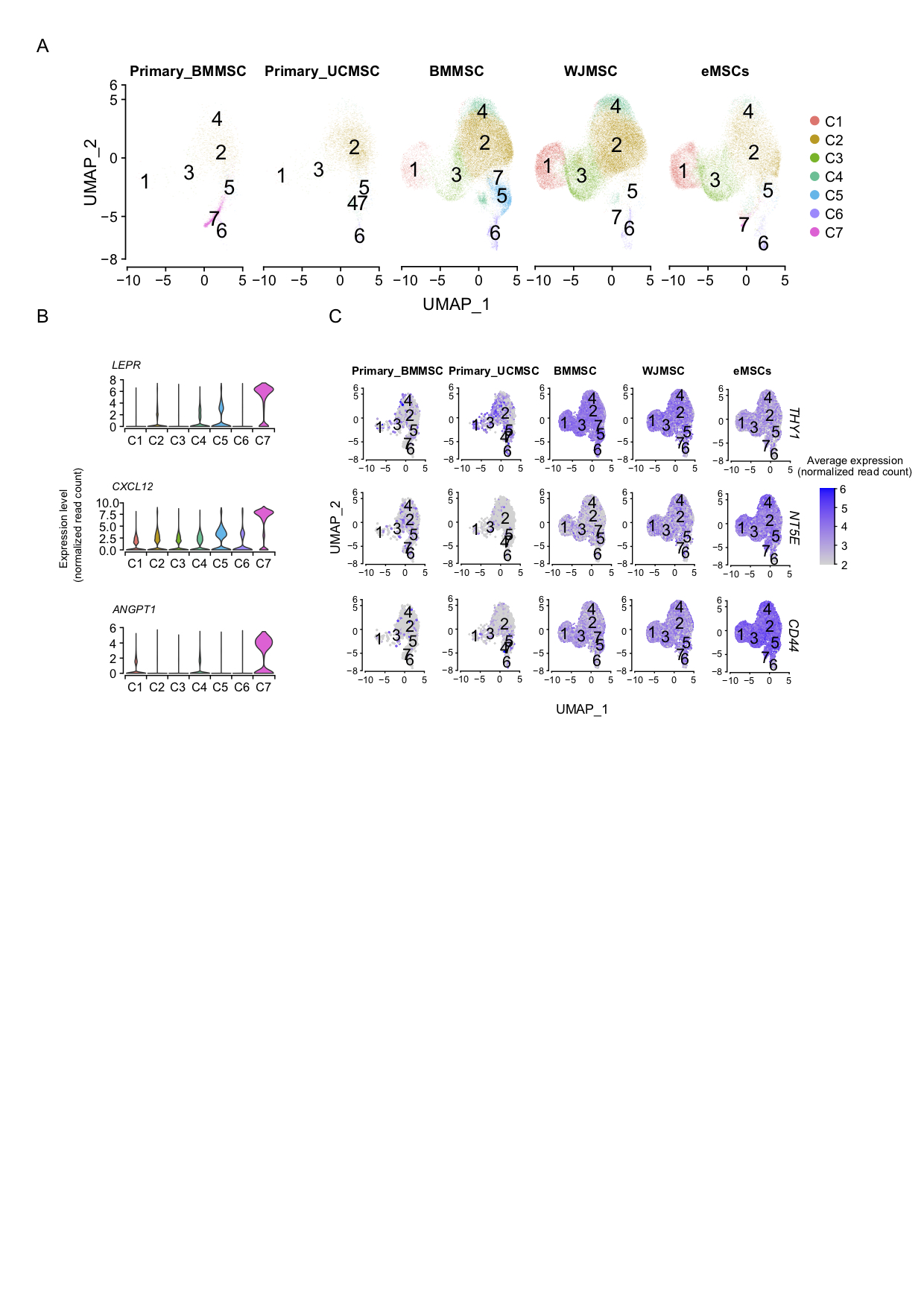
