## Supplemental Table 1 for "Single-cell Transcriptomic Analysis Reveals the Cellular Heterogeneity of Mesenchymal Stem Cells"

| | BMMSC1 | BMMSC2 | BMMSC3 | WJMSC1 | WJMSC2 | WJMSC3 |
|---|---|---|---|---|---|---|
| Abbreviation | HZX | CYX | GJY | GS-13 | GS-16 | cmo16 |
| Personal information | 2-year-old，cerebral palsy | 2-year-old，cerebral palsy | 2.5-year-old，cerebral palsy | Eutocia, no genetic disorders | Eutocia, no genetic disorders | 29-year-old，Eutocia, no genetic disorders |
| Sex | Female | Male | Female | Female | Female | Female |
| Passage | 6 | 6 | 7 | 6 | 6 | 6 |
| Cell viability | 89.89% | 90.11% | 81.99% | >90% | 95.73% | 92.99% |
| Aggregation rate | 10.39% | 19.23% | 12.73% | ≈20% | 38.92% | 8.36% |
| **Sequencing information** | | | | | | |
| Cell number | 13360 | 14071 | 12617 | 12756 | 12553 | 11154 |
| Mean reads per cell | 56759 | 52158 | 58706 | 66324 | 75602 | 68228 |
| Median genes per cell | 2896 | 2202 | 2937 | 4133 | 3838 | 4317 |
| Median UMI counts per cell | 11325 | 7263 | 11667 | 25658 | 19129 | 19652 |
| Total Genes Detected | 21613 | 21212 | 20969 | 22864 | 22319 | 22139 |
| Sequencing saturation | 60.80% | 59.80% | 57.80% | 40.40% | 56.60% | 51.30% |
| **Cells information after filtering** | | | | | | |
| Left cells | 10075 | 7499 | 10128 | 11655 | 11872 | 10067 |

**Table S1 Sample Information.**
